# Unleashing the potential of OPM-MEG to study event-related fields against low-frequency artifacts: the case of sentence processing

**DOI:** 10.1101/2025.03.14.643182

**Authors:** Xinchi Yu, Dongxue Zhang, Thomas Carey, Ernst Pöppel, Ellen Lau, Tilmann Sander

**Affiliations:** Program in Neuroscience and Cognitive Science, University of Maryland, College Park, MD, USA; Department of Linguistics, University of Maryland, College Park, MD, USA; Physikalisch-Technische Bundesanstalt, Berlin, Germany; Institute of Medical Psychology, Ludwig Maximilian University Munich, Munich, Germany; School of Psychological and Cognitive Sciences, Peking University, Beijing, China; Neurocognitive Image Lab, Shanghai International Studies University, Shanghai, China

**Keywords:** OPM-MEG, N400, sentence processing, DSS (denoising source separation)

## Abstract

OPM-MEG (optically pumped magnetometers-magnetoencephalography) offers unprecedented opportunity in its proximity to the brain and in mobility, outperforming other human MEG systems. However, movements induce low-frequency artifacts (up to ∼ 3 Hz) in this system, calling for pipelines that could efficiently reduce these low-frequency noises. Although a high-pass filter of e.g., 4 Hz may minimize these artifacts, it may also eliminate many event-related field (ERF) components, such as the N400 response in sentence processing studies. Moreover, as an emerging technology with expensive sensors, many labs are starting with an experimental setup with fewer sensors (e.g., ∼ 10), making it challenging to reject principal components based on visually inspecting the topographic field maps. In the current paper, we show that a combination of moderate high-pass filtering (1 Hz) and evoked-biased denoising source separation (evoked-biased DSS) can effectively reveal typical N400 deflections and effects (between nouns and verbs), with a nine-sensor OPM-MEG setup. Our current pipeline paves the way to studying ERFs with OPM-MEG in more budget-friendly setups with a small number of sensors.

## 1. Introduction

Magnetoencephalography (MEG) is a non-invasive neuroimaging technology with both high spatial and temporal resolution. MEG studies therefore offer important complementary evidence in the field of cognitive neuroscience, compared to e.g., EEG (electroencephalography; with good temporal resolution but bad spatial resolution) and fMRI (functional magnetic resonance imaging; with good spatial resolution but bad temporal resolution). MEG also possesses largely underexplored clinical potential including e.g., the diagnosis of concussion (Davenport et al., 2022; Lee & Huang, 2012) and dyslexia (Dikker et al., 2020; Salmelin, 2007; Tarkiainen et al., 2003). MEG sites existing for more than 10 years typically make use of SQUID-MEG systems (SQUID: superconducting quantum interference device), but recent years have seen an increasing transition to OPM-MEG systems (OPM: optically pumped magnetometers), which possess several properties that are more desirable than SQUID-MEG systems (Borna et al., 2017; Boto et al., 2017; Brickwedde et al., 2024; Brookes et al., 2022; Cheng et al., 2024; Kominis et al., 2003; Qin & Gao, 2021; Sander et al., 2012; Seymour et al., 2022). First, OPM-MEG systems do not require the constant liquid helium supply necessary for SQUIDs, which is expensive and is easily affected by helium shortages. Second, OPM sensors can be much closer to the scalp compared to SQUIDs, allowing for higher signal amplitudes (which is critical for detecting the faint magnetic signals from the brain, generally at the scale of femtoteslas [fT], 10^-15^ T). For example, in the current study the OPM sensors were attached to an EEG cap, which is much closer to neural sources in the brain compared to SQUIDs, which are encapsulated in a dewar that cannot fit flexibly to the scalp (in this case the minimum distance to cortical sources would be ∼ 2-4 cm; Andersen et al., 2020; Iivanainen et al., 2020). Third, most current OPM-MEG systems are much more mobile than SQUID-MEG systems, which may allow for mild movements during data collection. Mild movements are generally inevitable, especially for certain populations such as young children, positing challenges to the inclusion of more populations for SQUID-MEG systems. Moreover, a mobile setup allows for MEG recordings in more naturalistic settings.

However, what comes along with OPM-MEG’s mobility is a low-frequency artifact driven by the interaction between the magnetometers and the remnant field gradients present in all magnetically shielded rooms (Jazbinšek et al., 2022). Although participants are instructed not to move their head during experiments, it is impossible for their heads to be completely motionless due to e.g., breathing. Even seemingly small movements would drive considerable artifacts at the magnetometers, resulting in low-frequency artifacts in the data that are believed to be up to ∼ 3 Hz (Marhl et al., 2022; Seymour et al., 2022). This contrasts with canonical SQUID-MEG set-ups and some OPM-MEG designs (Alem et al., 2023), where the SQUID dewar or the OPM helmet are rigidly attached to the ground, and thus could only be affected when the ground moves relative to the Earth’s magnetic field (e.g., when someone is stomping on the ground close to the MEG scanner). Several previous studies would therefore apply a high-pass filter with a rather high cutoff of e.g., 4 Hz in order to ensure removal of low-frequency artifacts (Gialopsou et al., 2021; Jodko-Władzińska & Sander, 2024; Marhl et al., 2022; Seymour et al., 2021, 2022). While such a high high-pass filter (HPF) may not cause a problem if the goal is to examine the power of oscillatory bands above this filter, this practice may pose a problem for studying event-related fields (ERFs), as high-pass filters with a high cutoff are known to attenuate or eliminate ERP (event-related potential) amplitudes and ERP effects across conditions in EEG studies (Tanner et al., 2015; van Driel et al., 2021; G. Zhang et al., 2024a, 2024b). Moreover, since visual sentence processing studies usually present each segment (usually a word) for several hundreds of milliseconds, many ERF components may form some regularities in a lower frequency range slightly above 1 Hz (Ding et al., 2015; Lu et al., 2022), therefore a high-pass filter of 4 Hz would especially risk removing them (Seymour et al., 2022). On the other hand, some previous studies employed a moderate high-pass filter of e.g., 1 Hz, along with canonical preprocessing methods for SQUID-MEG data including ICA (Iivanainen et al., 2023). However, ICA may not be able remove the movement artifacts if they are statistically non-stationary. The fundamental challenge is that, while ICA procedures typically reject undesirable components related to e.g., eye blinks and heart beats through inspection of waveform and topographic field map (Puce & Hämäläinen, 2017), it is unclear what would be the canonical topography for low-frequency movement artifacts in OPM-MEG. Moreover, with experimental OPM-MEG setups with a handful of (e.g., ∼ 10) sensors, it is challenging to establish full topographic field maps in the first place (Iivanainen et al., 2023; Sander et al., 2020). Furthermore, the ability for ICA to single out “clean” artifact components appears to degrade as the number of sensors decrease (Klug & Gramann, 2021). However, the number of OPM sensors available is usually constrained by the nontrivial price of OPM sensors, so that many labs starting on OPM-MEG only possess a limited number of sensors. Therefore, a reasonable pipeline for OPM-MEG needs to be developed to analyze ERFs with the goal of (1) attenuating low-frequency movement artifacts, (2) retaining major ERF deflections and ERF effects, and (3) accommodating OPM-MEG setups with a smaller set of sensors (for which inspecting the topographic field map is challenging). Developing methods for setups with small numbers of OPMs will lead to more cost-effective research tools, as OPM sensors are still very expensive. Such pipelines would greatly unleash the potential of OPM-MEG to study ERFs, including in sentence processing experiments.

One existing denoising method, evoked-biased denoising source separation (evoked-biased DSS; de Cheveigné & Simon, 2008), may resolve this challenge, since this method does not rely on visual inspection of topographic field maps. Evoked-biased DSS decomposes data from sensors into PCA (principal component analysis) components, ranking from more time-locked to less time-locked. Assuming that movements are generally not time-locked to the stimulus, by keeping the first few components and excluding the rest of the components, we do not risk removing evoked responses much, while attenuating the effects from movement artifacts. However, evoked-biased DSS has not been tested on OPM-MEG data.

In the current OPM-MEG study, with a canonical visual sentence processing paradigm in Mandarin Chinese, we compared across preprocessing pipelines with various high-pass filtering thresholds and with/without evoked-biased DSS, leveraging the well-studied N400 response (and N400 effects across conditions) observed in SQUID-MEG and EEG (Lau et al., 2008, 2016). In SQUID-MEG, N400 manifests as a deflection around 350 ms which is particularly observable in the sensors above the left hemisphere (Pylkkänen et al., 2002, 2006; Stockall et al., 2004). Apart from the N400 deflection per se, we also capitalize on the difference in N400 amplitude between verbs and nouns, which has been observed in SQUID-MEG and EEG systems as well, across a variety of languages including Mandarin Chinese (our stimuli language; Federmeier et al., 2000; Khader et al., 2003; Tsigka et al., 2014; Xia et al., 2016; Xia & Peng, 2022). To date, few sentence processing experiments have been conducted with OPM-MEG. Although one recent study employed an auditory sentence processing paradigm (Wu et al., 2024), they primarily employed multivariate analysis methods, and so it was unclear whether the canonical univariate N400 deflection and N400 effects were preserved.

To preview, we observed that while a 4 Hz high-pass filter completely destroyed the N400 deflection and any N400 differences between verbs and nouns, a low (0.1 Hz) or moderate (1 Hz) high-pass filter alone retained too much low-frequency artifacts in ERF data. Combining a moderate high-pass filter (1 Hz) with evoked-biased DSS appeared to offer the best performance in attenuating low-frequency movement artifacts while retaining the N400 deflection and the N400 difference between verbs and nouns.

## 2. Method

### 2.1. Participants

11 native speakers of Mandarin Chinese participated in this study (5 females, age *M* = 25*, SD* = 3). Informed consent was obtained from all participants and they received monetary reimbursement for their participation. The procedures were approved by the IRB of University of Maryland.

### 2.2. Stimuli and procedure

The experiment was modeled after the EEG experiment of Yu et al. (2024), with more sentences and therefore more trials.

#### Stimuli

We first came up with two lists of 80 verbs and 80 nouns in Mandarin Chinese, which was a superset of the list used in Yu et al. (2024). The verbs consisted of 40 separable verbs and 40 inseparable verbs; the nouns consisted of 40 compound nouns and 40 simplex nouns. For their detailed syntactic properties, see Yu et al. (2024). Comparing the responses between the two types of verbs or the two types of nouns is beyond the purpose of the current paper. These verbs and nouns were all disyllabic, each corresponding to two Chinese characters (average number of strokes for each verb or noun item: verb: 16.2; noun: 15.3). Mean log frequency in the BCC corpus (Xun et al., 2016): verb: 3.9, noun: 3.5. Mean log frequency in the CCL corpus (Zhan et al., 2019): verb: 4.1, noun: 3.7.

Then we created 160 sentences based on the 160 verb and noun items. The verbs and nouns were embedded in sentences in a certain sentential structure respectively, demonstrated in Table 1. Two counterbalanced lists of sentences for the verbs were created with two counterbalanced list of people’s names. These two lists were administered to roughly half of the participants respectively. Another 60 filler sentences were adapted from another study (Liao & Lau, 2020), rendering in total 220 sentences for each participant.

**Table 1.** Example sentential stimuli for our experiment. The critical segment (i.e., the critical word) for our analysis is marked in bold.

|  |
| --- |
| **Nouns** |
| *Sentence structure:*<br>[因为] [**compound/simplex noun**] [是] [segment 4] [segment 5,] [所以/因此] [continuation.] |
| *English translation:*<br>[Because] [**compound/simplex noun**] [is] [segment 4] [segment 5,] [therefore/so] [continuation.] |
| **Example:**<br>因为/热狗/是/一种/高热量的/食物, /所以/不是很健康。<br>Because/**hotdog**/is/a.kind.of/high-calorie/food,/therefore/not.very.healthy.<br>“Because hotdog is a kind of high-calorie food, therefore it is not very healthy.” |
| **Verbs** |
| *Sentence structure:*<br>[Name] [想要/不想/很想] [**separable/inseparable verb,**] [所以/因此] [continuation.] |
| *English translation:*<br>[Name] [wants to/doesn't want to/really wants to] [**separable/inseparable verb,**] [so/therefore] [continuation.] |
| **Example:**<br>小范/很想/造反, /所以/他秘密组织了一支军队。<br>Fan[name]/really.wants.to/**rebel**,/therefore/he.gathered.an.army.secretly.<br>“Fan really wants to rebel, therefore he gathered an army secretly.” |

#### Procedure: sentence processing experiment

The procedure was also generally consistent with Yu et al. (2024). Participants sat ∼ 90 cm in front of the screen. Due to the setup of the projector in the lab, the screen had to be ∼ 30° tilted from the ground; the participants’ high performance suggested that the tilted screen did not affect sentence comprehension. 220 trials were run for each participant, with a different sentence presented in each trial. Each trial began with a fixation cross of 1000 ms with a 200 ms blank screen, followed by presentation of the segments of this sentence in a segment-by-segment (i.e., RSVP, rapid serial visual presentation) manner, with a presentation duration for 600 ms for each segment, and a 200 ms interval between segments, for a total of 800 ms stimulus-onset asynchrony (Figure 1). The segments were presented in white characters against a black background, in ∼ 57 pt, Yahei UI font, at the center of the screen. In ∼ 1/3 of the trials (76 out of 220 trials), the last segment of the sentence (i.e., the continuation) was colored in green and the subjects were instructed to respond to whether this continuation was congruent in this sentence, by pressing the left (congruent) or the right button (incongruent) on a response pad (the responses did not have a time-limit). In the other ∼ 2/3 of the trials, the last segment was presented on the screen in white briefly for 1000 ms, and subjects did not need to respond. The inter-trial intervals were self-paced; subjects pressed any button when they were ready to start the next trial. The participants completed a SQUID-MEG session with similar materials prior to the OPM-MEG visit. Since the two visits were interleaved by on average 10.5 (range 5-18) days, the SQUID-MEG session would unlikely affect the OPM-MEG session.

**Figure 1.**
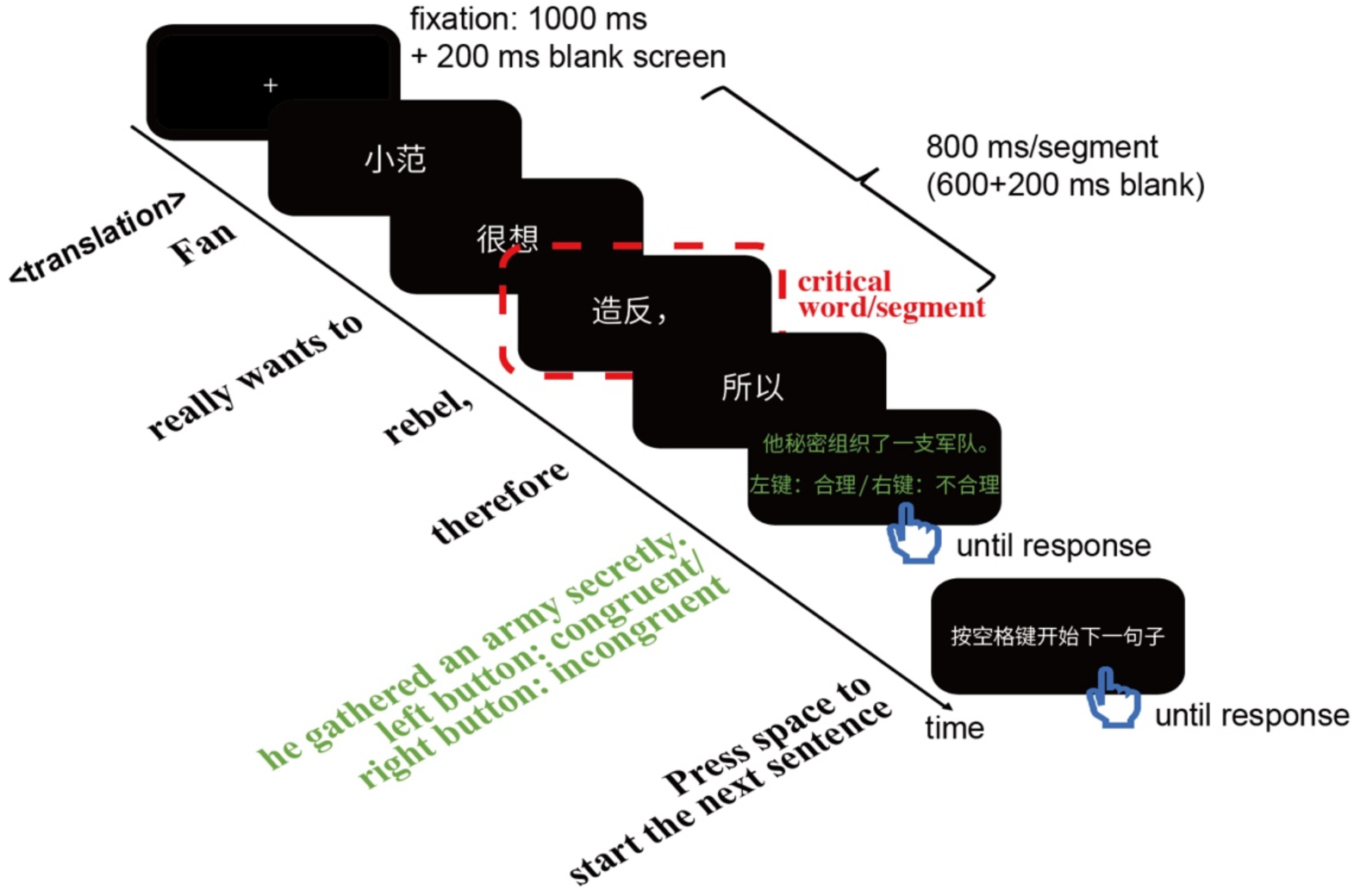
Illustration of a trial of our sentence processing study. The English translation of the stimuli presented in Chinese is below the arrow.

### 2.3. Behavioral data analysis

Behavioral accuracy for each participant was calculated for the trials with a continuation judgement task, as the proportion of correct responses.

### 2.4. OPM Setup

The OPM measurements were conducted in a two-layered Ak3b magnetically shielded room (Vacuumschmelze GmbH & Co. KG, Hanau, Germany). Nine OPM magnetometer sensors (QZFM generation 2, QuSpin Inc., Louisville, Colorado, USA) were positioned across the scalp using a modified EEG cap (easycap 21-channel 10-20 cap). The cap had clip in OPM holders attached to the fabric (similar to the approach in https://quspin.com/experimental-meg-cap/). Such fabric caps modified for OPM usage have been used for clinical works as well (Feys et al., 2022). OPMs were placed especially on the left hemisphere, at T8, F4, Fpz, F3, F7, C3, T7, P7, O1 and O2 (Figure 2). Because the easycap 21-channel cap doesn’t have an Fpz location, we created one at the appropriate location between Fp1 and Fp2. Each sensor recorded from two axes (z and y: radial head field and tangential field). The direction of the y axis was not consistent across all subjects due to rotational uncertainties, but was constant within each subject. In the current paper, we only analyze the z axis (radial direction relative to the head), which is in line with the direction recorded in most existing SQUID-MEG devices. Although SQUID devices mainly consist of gradiometers, these gradiometers are designed to be sensitive to radial magnetic field information.

**Figure 2.**
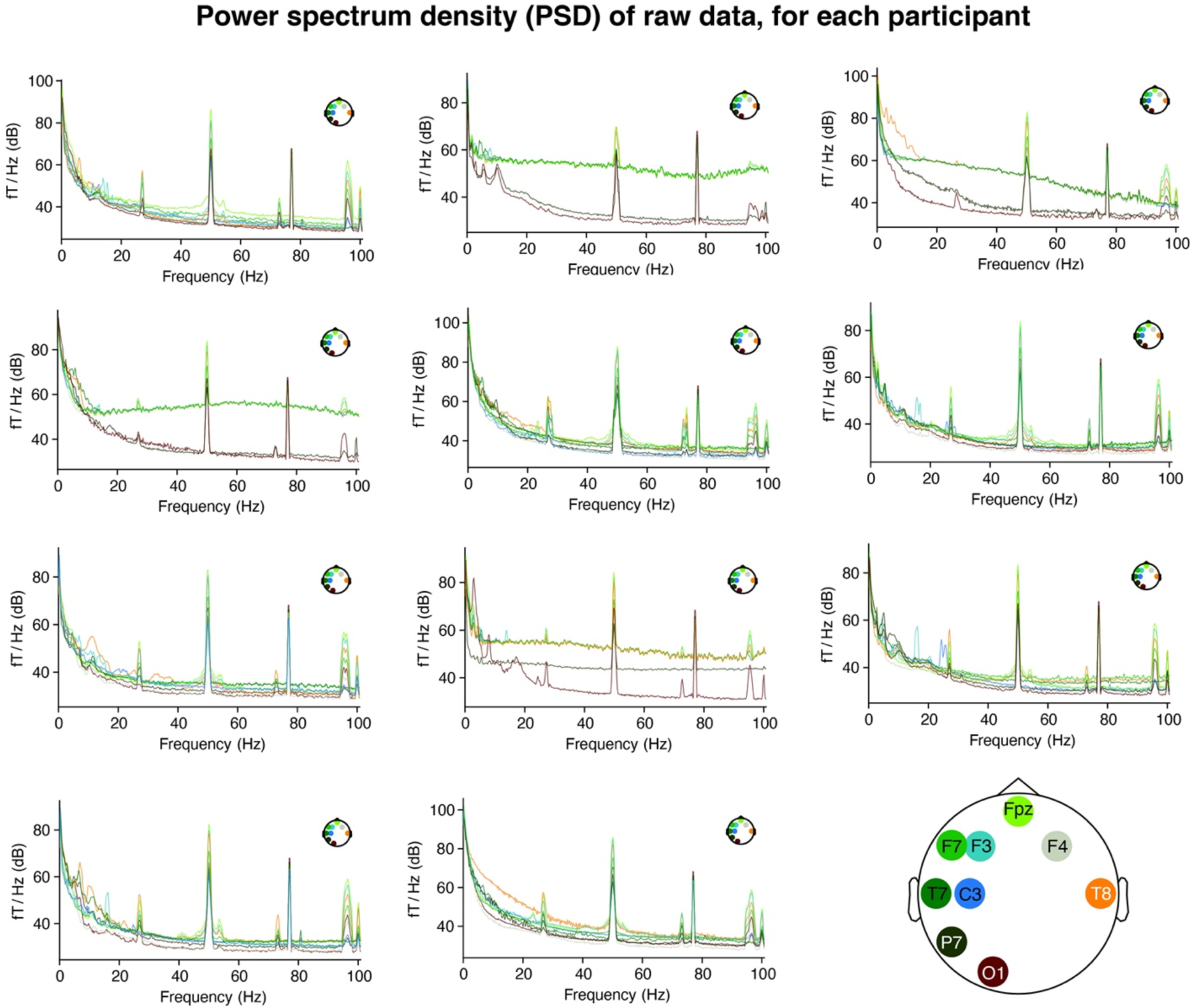
Power spectrum density (PSD) of raw OPM-MEG data for each of the 11 participants. The OPM sensors were attached to an EEG cap, whose layout is illustrated in the bottom right corner. Consistent with prior OPM-MEG studies, we observed two spectral peaks around 27 Hz (cause yet unknown) and 50 Hz (line noise) across most participants.

### 2.5. OPM data analysis

OPM data analyses was conducted with customized code in Python and MATLAB, with MNE-Python (Gramfort et al., 2013). Statistical tests (paired t-test and repeated-measures ANOVA) were administered in JASP 0.18.1 (JASP Team, 2023).

#### Preprocessing

Upon inspecting the power spectrum (Figure 2) for individual participants, there were generally a 50 Hz component as well as a ∼ 27 Hz component. The 50 Hz component was most likely line noise. A 20-30 Hz component is generally present in other OPM systems as well (Alem et al., 2023; Seymour et al., 2022; X. Zhang et al., 2020) with a yet-unclear source. Therefore, we applied notch filters at 50 and 27 Hz (transitional bandwidth = 4 Hz) to remove these components. Then we tested six different treatments: (1) Data was passed with a 0.1-40 Hz IIR (infinite impulse response) filter (i.e., high-pass filter, HPF = 0.1 Hz) only, (2) Data was passed with a 0.1-40 Hz IIR filter, then an evoked-biased denoising source separation (evoked-biased DSS), (3) a 1-40 Hz IIR filter only, (4) a 1-40 Hz IIR filter followed by evoked-biased DSS, (5) a 4-40 Hz IIR filter only, (6) a 4-40 Hz IIR filter followed by evoked-biased DSS. The other parameters for the IIR filters were determined by default algorithms in MNE-Python.

The procedure of evoked-biased DSS (de Cheveigné & Simon, 2008) is as follows. Evoked-biased DSS transforms normalized data from 9 sensors into 9 components, ranking from more stimulus-evoked-like to more noise-like. By keeping the first few components and excluding the rest of the components, we do not risk removing evoked responses much, while attenuating the effects from movement artifacts. The final epoch size that we wanted to have is −200:800 ms, time-locked to critical word onset. Each epoch was baselined based on the −200:0 ms baseline. In order to reduce the influence of the baseline period in DSS (note: in the current paper, “DSS” refers to evoked-biased DSS), we ran DSS on longer epochs of −200:1600 ms (Kim et al., 2020, 2022). Evoked-biased DSS was conducted with all epochs across the four conditions (epoch data were normalized for each sensor prior to DSS), resulting in 9 DSS components. We projected the first two DSS components back to the sensor space to create corrected epochs for analysis.

ERFs (event-related fields) were computed as the median of the epochs for each participant and for each condition respectively: separable verb, inseparable verb, compound noun, simplex noun, aggregated verb condition across separable and inseparable verbs, as well as aggregated noun condition across compound and simplex nouns. Because the mean is more subject to the influence of outlier data points, computing mean ERFs would require us to apply an additional outlier cutoff processing step which would likely result in different numbers of excluded trials across preprocessing pipelines. Median ERFs are less sensitive to outliers, therefore computing the ERFs with the median mitigates the need for applying a cutoff.

#### Calculation of cross-trial variance for ERFs

As a means of quantifying overall noise level in the results of each pipeline, we computed a cross-trial variance measure for the ERFs. For each pipeline, for each participant and at each time point, we first calculated *one* variance value across all verb epochs and all noun epochs respectively. This variance value was calculated in the following way. First, we calculated the standard deviation at each sensor. Then, we averaged the standard deviations across all 9 sensors. After that, for each time point, we averaged the standard deviations across all participants, for the noun epochs and verb epochs respectively. This way, for noun epochs and verb epochs respectively, we derived an 1D array across time, with one averaged variance value at each time point.

#### Calculation of RMS (root mean square)

To determine whether the pipelines adequately captured the typical peaks in the ERF, we used an RMS measure. We calculated the RMS from ERFs that were first averaged across the verb and noun conditions, then averaged across participants. RMS was calculated as the root mean square across all 9 sensors.

#### Calculation of N400 amplitude

As described below, ERFs collapsed across conditions indicated a clear N400 peak between 300-400ms that was maximal at sensor F7. Therefore, we calculated N400 amplitude for each condition and for each participant at this sensor as the mean response amplitude across the 300-400 ms time window upon segment onset.

## 3. Results

### 3.1. Behavioral results

#### Accuracy

Participants generally achieved ceiling accuracy (range 99%-100%) in judging the semantic congruity of sentence continuations. Therefore, all trials were included in the MEG analysis.

### 3.2. MEG results

In this section we evaluate the processing pipelines based on several qualitative and quantitative aspects.

#### Shape of ERF

In pipelines utilizing a 1 Hz high-pass filter (HPF), we were able to identify a clear deflection around 400 ms in left centro-frontal sensors, including F7, T7 and C3 (Figure 3). The deflection had the highest amplitude at F7, from which we calculate the amplitude of N400 responses for further analysis. This deflection was faintly identifiable when using a 0.1 Hz HPF, and was practically invisible when applying a 4 Hz HPF. The same observation generally holds when inspecting individual participants’ data (Figure 4). Notably, the N400 deflection we observed was at the magnitude of hundreds of fT, which is higher compared to the typical magnitude observed in SQUID-MEG systems, at the magnitude of tens of fT (Lau et al., 2016; Lau & Nguyen, 2015), as expected given the much closer proximity of OPM-MEG sensors to the underlying neural generators. Additionally, this difference in magnitude may also be relevant to the fact that, while OPM sensors are magnetometers, SQUIDs are often designed in a gradiometer configuration.

**Figure 3.**
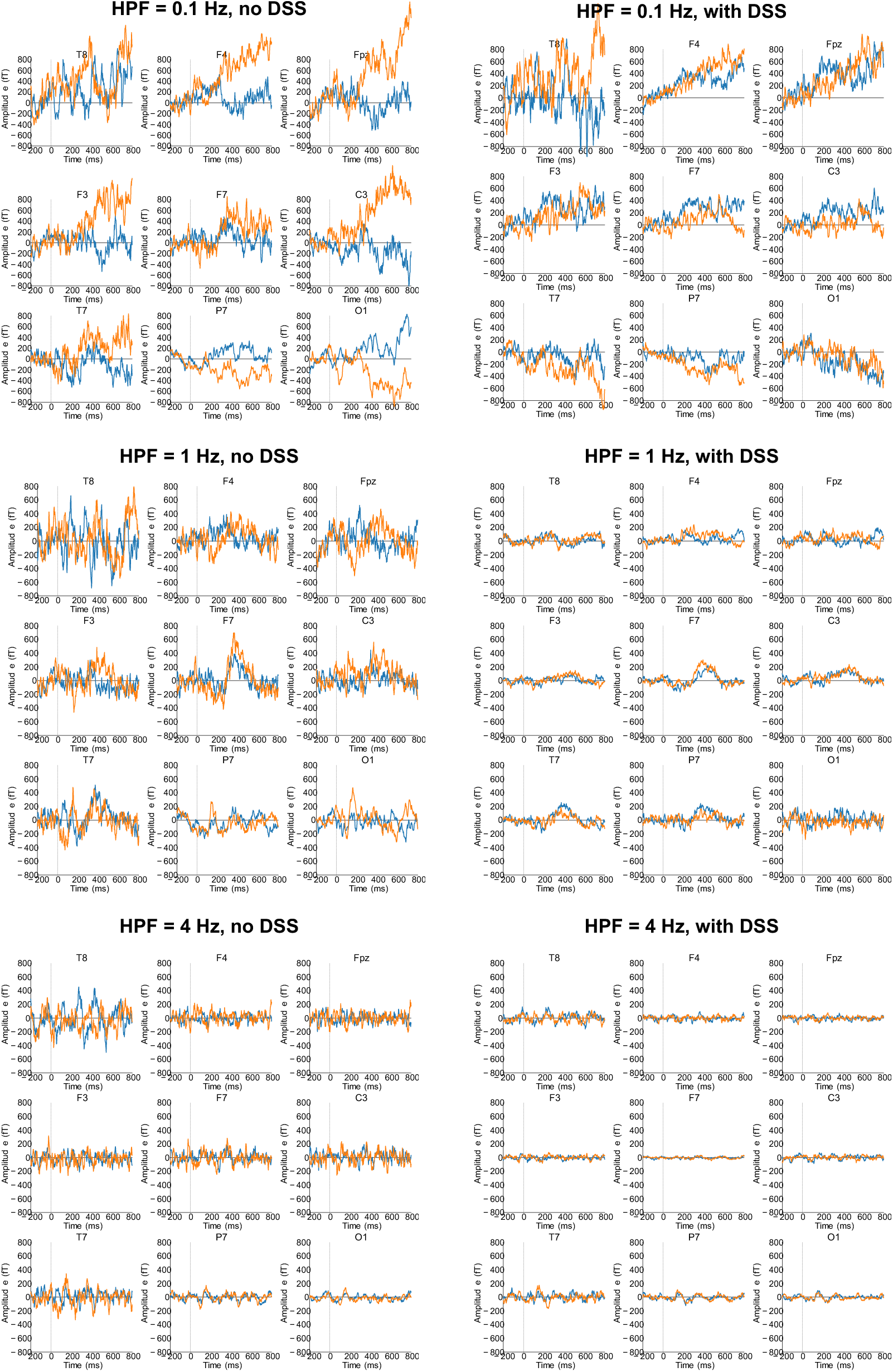
Grand average ERFs across all participants (blue line: verb; orange line: noun). ERF: event-related field.

**Figure 4.**
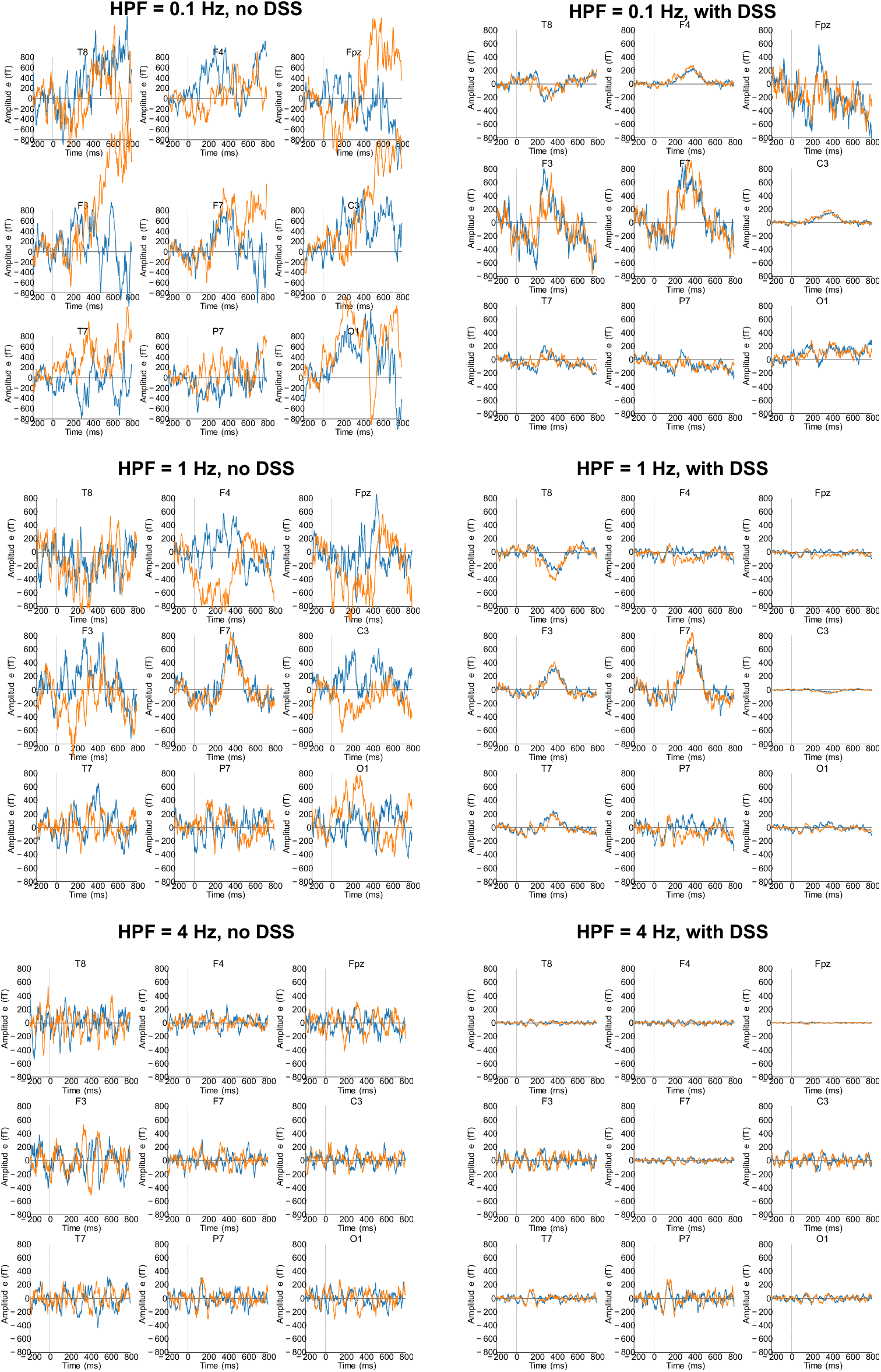
Median ERFs for noun and verb epochs in one participant (blue line: verb; orange line: noun). ERF: event-related field. The figures for every individual participant are shared openly on OSF.

#### Cross-trial variance for ERFs

Across the entire epoch, the pipeline with HPF = 4 Hz, with DSS had the lowest cross-trial variance, which holds for both verb trials and noun trials (Figure 5). This was followed by the pipeline with HPF = 1Hz, with DSS. The pipeline with HPF = 0.1 Hz without DSS had the highest variance among the pipelines.

**Figure 5.**
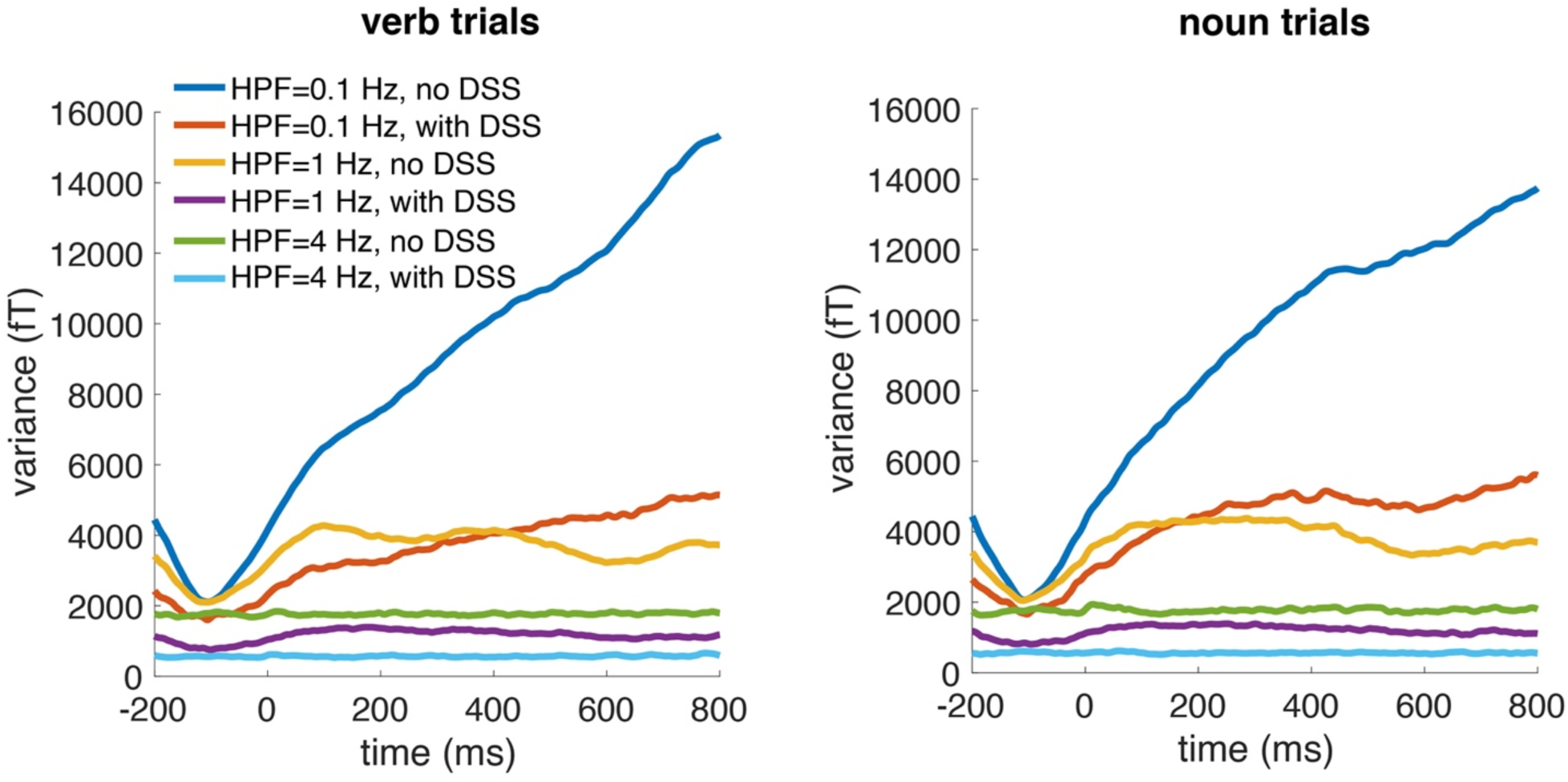
Cross-trial variance for each pipeline among verb trials (left) and noun trials (right), averaged across all participants. HPF: high-pass filter; DSS: denoising source separation.

#### Shape of RMS

Previous MEG sentence or single-word processing studies tend to observe a ∼ 400 ms peak, along with earlier peak(s) around ∼ 200 ms in RMS (Assadollahi & Pulvermüller, 2003; Maess et al., 2016; Tsigka et al., 2014). The ∼ 200 ms peak(s) likely reflect visual-orthographic processing (Gwilliams et al., 2016; Salmelin, 2007). The ∼ 400 ms peak or bump was clearly visible when we employed a 1 Hz HPF, while being faint or invisible for 0.1 or 4 Hz HPFs (Figure 6). A peak ∼ 200 ms can only be clearly visually identified when a 1 Hz HPF is combined with DSS, but not when only applying a 1 Hz HPF.

**Figure 6.**
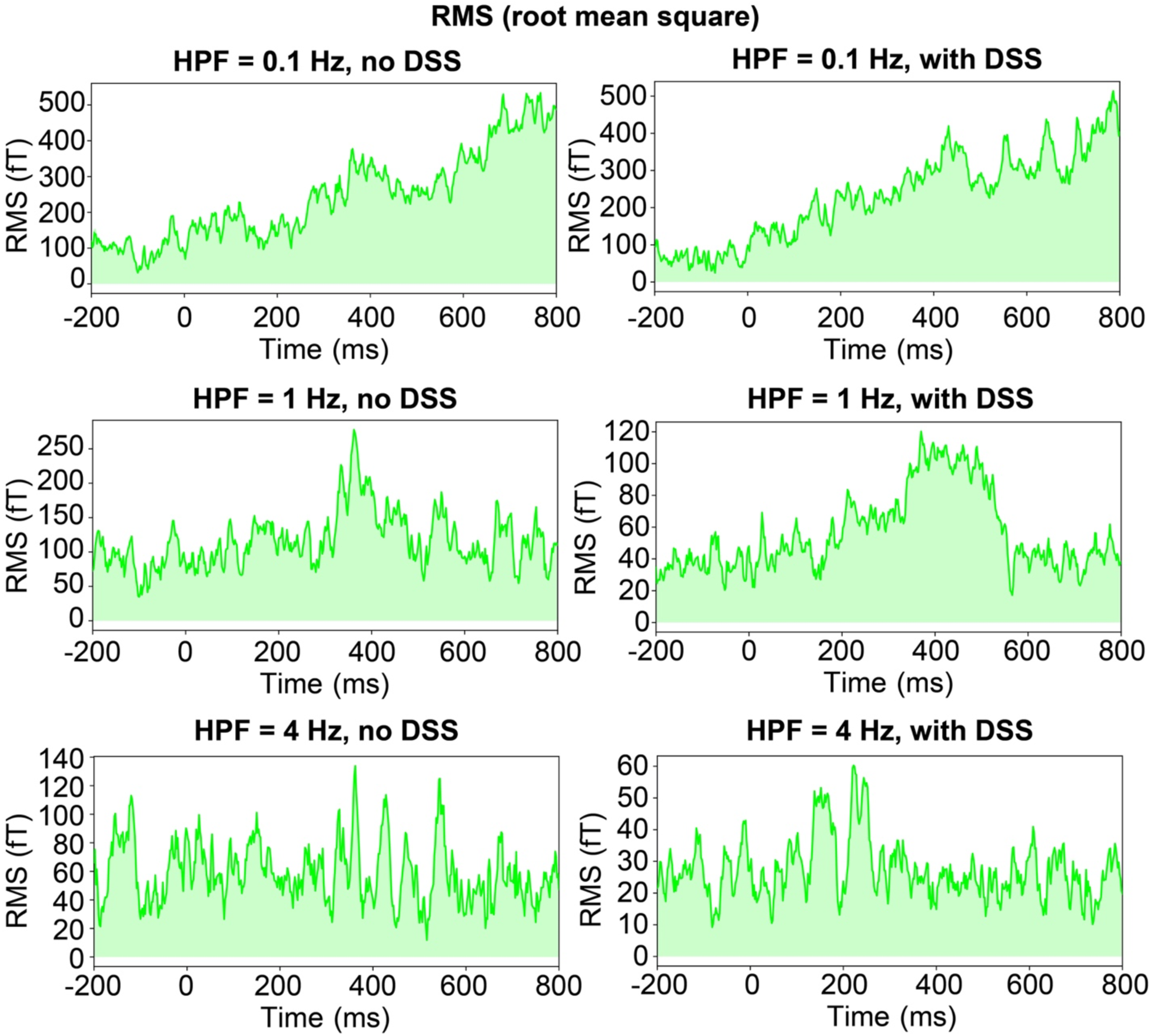
RMS (root mean square) across all electrodes, based on the ERFs averaged across verbs and nouns. Only when combining a 1 Hz HPF and DSS are we able to clearly visually identify peaks around 200 ms and 400 ms. ERF: event-related field; HPF: high-pass filter; DSS: denoising source separation.

#### N400 effect between verbs and nouns

Paired t-test (two-tailed) only revealed a significant difference between the N400 amplitudes of verbs and nouns for the pipeline with HPF = 1 Hz and DSS, *t*(10) = −2.93, *p* = 0.015, Cohen’s d = −0.89 (Figure 7). This difference was not significantly different for the other pipelines, although the effects were generally in the same direction. HPF = 0.1 Hz without DSS: *t*(10) = −0.36, *p* = 0.73, Cohen’s d = −0.11; HPF = 0.1 Hz with DSS: *t*(10) = 1.48, *p* = 0.17, Cohen’s d = 0.45; HPF = 1 Hz without DSS: *t*(10) = −1.67, *p* = 0.13, Cohen’s d = −0.50; HPF = 4 Hz without DSS: *t*(10) = −1.27, *p* = 0.23, Cohen’s d = −0.38; HPF = 4 Hz with DSS: *t*(10) = −0.83, *p* = 0.43, Cohen’s d = −0.25. Repeated-measures ANOVA across the four conditions (the two verb conditions and the two noun conditions) showed similar results (see Supplementary Materials).

**Figure 7.**
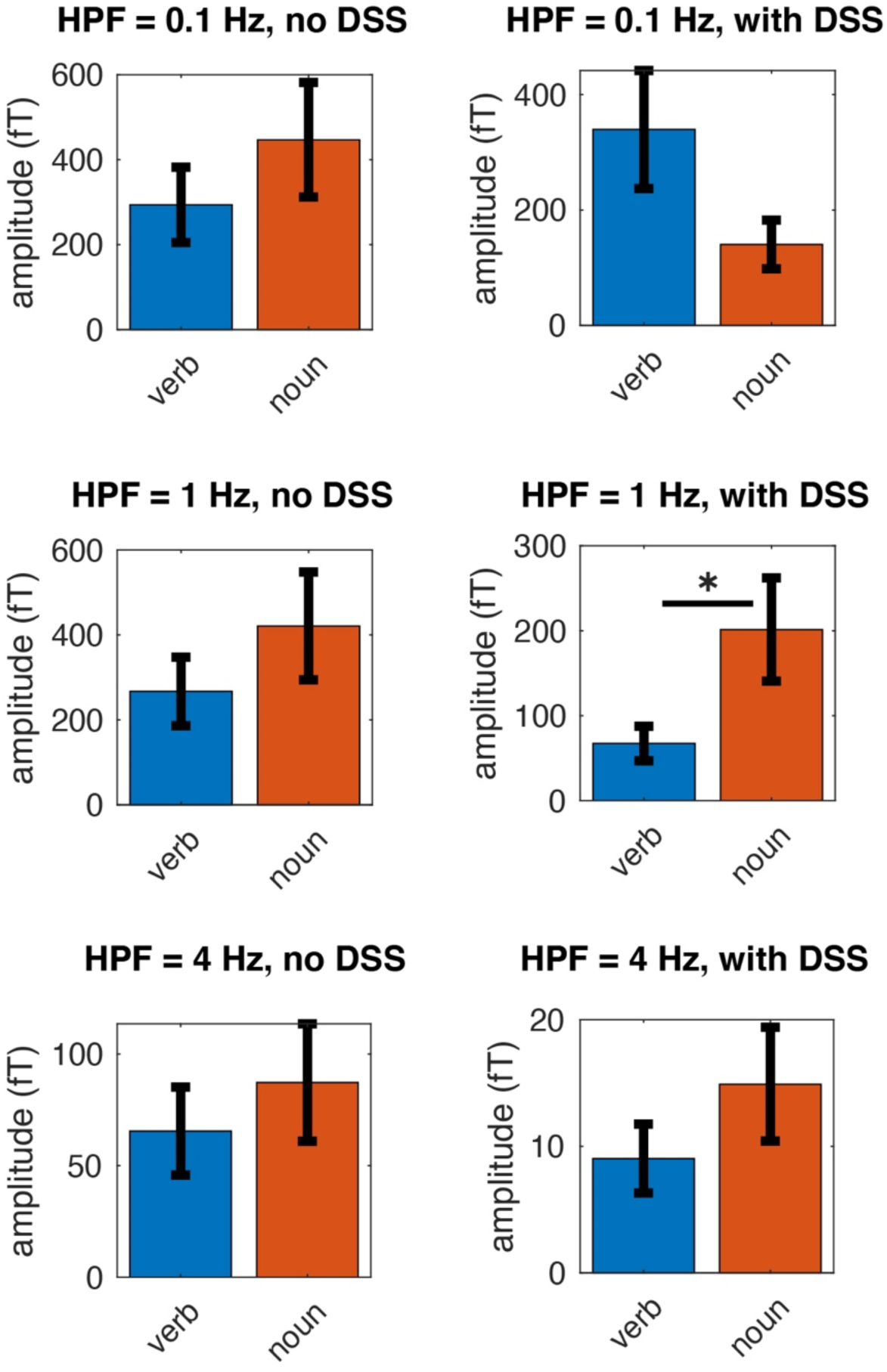
Comparison of N400 response (300-400 ms) amplitude at sensor F7 across verbs and nouns. Only when an HPF (high-pass filter) of 1 Hz was combined with DSS, were we able to observe a significant difference across verbs and nouns. Error bars stand for standard error. *: *p* < 0.05. HPF: high-pass filter; DSS: denoising source separation.

## 4. Conclusion

In the current paper, by comparing different preprocessing pipelines for OPM-MEG data, we demonstrated that combining a moderate high-pass filter (HPF) of 1 Hz with evoked-biased DSS preserves the N400 deflection and allowed us to successfully replicate prior observations of an N400 difference between nouns and verbs. Moreover, this pipeline revealed canonical RMS peaks, and had a relatively low cross-trial variance among the pipelines. This pipeline thus unleashes the potential of OPM-MEG to study ERFs against low-frequency artifacts, especially for OPM-MEG setups with fewer sensors, where methods like ICA would be inapplicable. This can allow a beneficial ‘convergent methods’ approach for labs to obtain high-quality MEG data in closer proximity to the neural generators than traditional systems, even in the face of current constraints on the number of OPM sensors.

## 5. Future directions

### 5.1. On applying higher high-pass filters

Although a 4 Hz HPF largely destroyed the N400 deflection and N400 effects in our study, such higher HPFs may still be applicable in studying some other ERFs. For example, the auditory evoked responses of M100, which is a peak ∼ 100 ms upon auditory stimuli onset, appeared to survive such higher HPFs in previous studies (Marhl et al., 2022). Notably, in Marhl et al. (2022), auditory pure tones were presented at a duration of 500 ms, with intervals of 1200 ms between the tones. Thus, the stimuli presentation would form regularities below the 4 Hz HPF. Nevertheless, M100 was observed in that study. This is not surprising since M100 is a transient response that only lasted for ∼ 50 ms, compared to our N400 deflections, which appeared to extend for ∼ 300 ms. Exactly when higher HPFs can/cannot be used to study which ERF component remains to be further elucidated in future studies, through empirical investigations and simulations.

### 5.2. Other pipelines

Beyond our current pipeline, some other pipelines may also achieve satisfactory performances, and future studies could work on comparing across these pipelines. Of note, HFC (homogeneous field correction) may be a promising method for this purpose (Tierney et al., 2021). We did not perform HFC for our current data because it requires the sensor orientation angles, which we did not record in this study. Future studies could compare across our current pipeline and HFC for the metrics that we used in the current work, or even combine them together.

### 5.3. Exploring OPM-MEG recording with a limited number of sensors

In the current study, we demonstrated a pipeline that accommodates OPM-MEG systems with a limited number of sensors (in our case, nine sensors), and obtained meaningful ERF responses in a sentence processing study. In practice, the number of sensors may not only be constrained by financial costs, but also by preparation time. For example, attaching more OPM sensors to the cap increases setup time, which may be a limiting factor in certain clinical contexts where shorter preparation times are preferred. Notably, previous research has shown that as few as four dual-axis OPM sensors can be sufficient for a brain-computer interface for motor imagery decoding (Tan et al., 2025). This, along with our findings, suggests that even with a reduced sensor count, OPM-MEG can still yield meaningful results. Taken together, our study marks one of the initial steps in validating OPM-MEG systems with fewer sensors, paving the way for more cost-effective solutions which would enable wider clinical usage of OPM-MEG.

## Data and Code Availability

Data, stimuli and processing scripts for the current study will be openly shared on the Open Science Framework upon publication.

## Author contributions

**Xinchi Yu**: conceptualization, methodology, formal analysis, investigation, data curation, visualization, funding acquisition, writing – original draft, writing – review & editing

**Dongxue Zhang**: software, investigation, writing – review & editing

**Thomas Carey**: investigation, writing – review & editing

**Ernst Pöppel**: resources, supervision, writing – review & editing

**Ellen Lau**: conceptualization, supervision, writing – review & editing

**Tilmann Sander**: methodology, resources, supervision, writing – review & editing

## Acknowledgements

We would like to thank Jonathan Simon for discussions.

## References

Alem, O., Hughes, K. J., Buard, I., Cheung, T. P., Maydew, T., Griesshammer, A., Holloway, K., Park, A., Lechuga, V., Coolidge, C., Gerginov, M., Quigg, E., Seames, A., Kronberg, E., Teale, P., & Knappe, S. (2023). An integrated full-head OPM-MEG system based on 128 zero-field sensors. Frontiers in Neuroscience, 17(June), 1–8. 10.3389/fnins.2023.1190310

Andersen, L. M., Pfeiffer, C., Ruffieux, S., Riaz, B., Winkler, D., Schneiderman, J. F., & Lundqvist, D. (2020). On-scalp MEG SQUIDs are sensitive to early somatosensory activity unseen by conventional MEG. NeuroImage, 221(July). 10.1016/j.neuroimage.2020.117157

Assadollahi, R., & Pulvermüller, F. (2003). Early influences of word length and frequency: A group study using MEG. NeuroReport, 14(8), 1183–1187. 10.1097/00001756-200306110-00016

Borna, A., Carter, T. R., Goldberg, J. D., Colombo, A. P., Jau, Y. Y., Berry, C., McKay, J., Stephen, J., Weisend, M., & Schwindt, P. D. D. (2017). A 20-channel magnetoencephalography system based on optically pumped magnetometers. Physics in Medicine and Biology, 62(23), 8909–8923. 10.1088/1361-6560/aa93d1

Boto, E., Meyer, S. S., Shah, V., Alem, O., Knappe, S., Kruger, P., Fromhold, T. M., Lim, M., Glover, P. M., Morris, P. G., Bowtell, R., Barnes, G. R., & Brookes, M. J. (2017). A new generation of magnetoencephalography: Room temperature measurements using optically-pumped magnetometers. NeuroImage, 149(January), 404–414. 10.1016/j.neuroimage.2017.01.034

Brickwedde, M., Anders, P., Kühn, A. A., Lofredi, R., Holtkamp, M., Kaindl, A. M., Grent-‘t-Jong, T., Krüger, P., Sander, T., & Uhlhaas, P. J. (2024). Applications of OPM-MEG for translational neuroscience: a perspective. Translational Psychiatry, 14(1), 1–12. 10.1038/s41398-024-03047-y

Brookes, M. J., Leggett, J., Rea, M., Hill, R. M., Holmes, N., Boto, E., & Bowtell, R. (2022). Magnetoencephalography with optically pumped magnetometers (OPM-MEG): the next generation of functional neuroimaging. Trends in Neurosciences, 45(8), 621–634. 10.1016/j.tins.2022.05.008

Cheng, H., He, K., Li, C., Ma, X., Zheng, F., Xu, W., Liao, P., Yang, R., Li, D., Qin, L., Na, S., Lyu, B., & Gao, J. H. (2024). Wireless optically pumped magnetometer MEG. NeuroImage, 300(April), 120864. 10.1016/j.neuroimage.2024.120864

Davenport, E. M., Urban, J. E., Vaughan, C., DeSimone, J. C., Wagner, B., Espeland, M. A., Powers, A. K., Whitlow, C. T., Stitzel, J. D., & Maldjian, J. A. (2022). MEG measured delta waves increase in adolescents after concussion. Brain and Behavior, 12(9), 1–9. 10.1002/brb3.2720

de Cheveigné, A., & Simon, J. Z. (2008). Denoising based on spatial filtering. Journal of Neuroscience Methods, 171(2), 331–339. 10.1016/j.jneumeth.2008.03.015

Dikker, S., Assaneo, M. F., Gwilliams, L., Wang, L., & Kösem, A. (2020). Magnetoencephalography and Language. Neuroimaging Clinics of North America, 30(2), 229–238. 10.1016/j.nic.2020.01.004

Ding, N., Melloni, L., Zhang, H., Tian, X., & Poeppel, D. (2015). Cortical tracking of hierarchical linguistic structures in connected speech. Nature Neuroscience, 19(1), 158–164. 10.1038/nn.4186

Federmeier, K. D., Sega, J. B., Lombrozo, T., & Kutas, M. (2000). Brain responses to nouns, verbs and class-ambiguous words in context. Brain, 123(12), 2552–2566. 10.1093/brain/123.12.2552

Feys, O., Corvilain, P., Aeby, A., Sculier, C., Christiaens, F., Holmes, N., Brookes, M., Goldman, S., Wens, V., & De Tiège, X. (2022). On-Scalp Optically Pumped Magnetometers versus Cryogenic Magnetoencephalography for Diagnostic Evaluation of Epilepsy in School-aged Children. Radiology, 304(2), 429–434. 10.1148/radiol.212453

Gialopsou, A., Abel, C., James, T. M., Coussens, T., Bason, M. G., Puddy, R., Lorenzo, F. Di, Rolfs, K., Voigt, J., Sander, T., Cercignani, M., & Krüger, P. (2021). Improved spatio - temporal measurements of visually evoked fields using optically - pumped magnetometers. Scientific Reports, 1–11. 10.1038/s41598-021-01854-7

Gramfort, A., Luessi, M., Larson, E., Engemann, D. A., Strohmeier, D., Brodbeck, C., Goj, R., Jas, M., Brooks, T., Parkkonen, L., & Hämäläinen, M. (2013). MEG and EEG data analysis with MNE-Python. Frontiers in Neuroscience, 7(7 DEC), 1–13. 10.3389/fnins.2013.00267

Gwilliams, L., Lewis, G. A., & Marantz, A. (2016). Functional characterisation of letter-specific responses in time, space and current polarity using magnetoencephalography. NeuroImage, 132, 320–333. 10.1016/j.neuroimage.2016.02.057

Iivanainen, J., Carter, T. R., Trumbo, M. C. S., McKay, J., Taulu, S., Wang, J., Stephen, J. M., Schwindt, P. D. D., & Borna, A. (2023). Single-trial classification of evoked responses to auditory tones using OPM- and SQUID-MEG. Journal of Neural Engineering, 20(5), 056032. 10.1088/1741-2552/acfcd9

Iivanainen, J., Zetter, R., & Parkkonen, L. (2020). Potential of on-scalp MEG: Robust detection of human visual gamma-band responses. Human Brain Mapping, 41(1), 150–161. 10.1002/hbm.24795

JASP Team. (2023). JASP 0.18.1.

Jazbinšek, V., Marhl, U., & Sander, T. (2022). SERF-OPM Usability for MEG in Two-Layer-Shielded Rooms. In Flexible High Performance Magnetic Field Sensors (pp. 179–193). Springer International Publishing. 10.1007/978-3-031-05363-4_10

Jodko-Władzińska, A., & Sander, T. (2024). Emotionally Charged Visually Evoked Magnetic Fields. Acta Physica Polonica A, 146(4), 521–525. 10.12693/APhysPolA.146.521

Khader, P., Scherag, A., Streb, J., & Rösler, F. (2003). Differences between noun and verb processing in a minimal phrase context: A semantic priming study using event-related brain potentials. Cognitive Brain Research, 17(2), 293–313. 10.1016/S0926-6410(03)00130-7

Kim, S. G., Overath, T., Sedley, W., Kumar, S., Teki, S., Kikuchi, Y., Patterson, R., & Griffiths, T. D. (2022). MEG correlates of temporal regularity relevant to pitch perception in human auditory cortex. NeuroImage, 249(January). 10.1016/j.neuroimage.2022.118879

Kim, S. G., Poeppel, D., & Overath, T. (2020). Modulation change detection in human auditory cortex: evidence for asymmetric, nonlinear edge detection. The European Journal of Neuroscience, January, 1–16. 10.1111/ejn.14707

Klug, M., & Gramann, K. (2021). Identifying key factors for improving ICA-based decomposition of EEG data in mobile and stationary experiments. European Journal of Neuroscience, 54(12), 8406–8420. 10.1111/ejn.14992

Kominis, I. K., Kornack, T. W., Allred, J. C., & Romalis, M. V. (2003). A subfemtotesla multichannel atomic magnetometer. Nature, 422(6932), 596–599. 10.1038/nature01484

Lau, E., & Nguyen, E. (2015). The role of temporal predictability in semantic expectation: An MEG investigation. Cortex, 68, 8–19. 10.1016/j.cortex.2015.02.022

Lau, E., Phillips, C., & Poeppel, D. (2008). A cortical network for semantics: (De)constructing the N400. Nature Reviews Neuroscience, 9(12), 920–933. 10.1038/nrn2532

Lau, E., Weber, K., Gramfort, A., Hämäläinen, M. S., & Kuperberg, G. R. (2016). Spatiotemporal Signatures of Lexical-Semantic Prediction. Cerebral Cortex, 26(4), 1377–1387. 10.1093/cercor/bhu219

Lee, R. R., & Huang, M. (2012). Magnetoencephalography in the diagnosis of concussion. *Concussion*, November, 94–111. 10.1159/000358768

Liao, C. H., & Lau, E. (2020). Enough time to get results? An ERP investigation of prediction with complex events. Language, Cognition and Neuroscience, 35(9), 1162–1182. 10.1080/23273798.2020.1733626

Lu, Y., Jin, P., Pan, X., & Ding, N. (2022). Delta-band neural activity primarily tracks sentences instead of semantic properties of words. NeuroImage, 118979. 10.1016/j.neuroimage.2022.118979

Maess, B., Mamashli, F., Obleser, J., Helle, L., & Friederici, A. D. (2016). Prediction signatures in the brain: Semantic pre-activation during language comprehension. Frontiers in Human Neuroscience, 10(NOV2016), 1–11. 10.3389/fnhum.2016.00591

Marhl, U., Jodko-Władzińska, A., Brühl, R., Sander, T., & Jazbinšek, V. (2022). Transforming and comparing data between standard SQUID and OPM-MEG systems. PLoS ONE, 17(1 January), 1–22. 10.1371/journal.pone.0262669

Puce, A., & Hämäläinen, M. S. (2017). A review of issues related to data acquisition and analysis in EEG/MEG studies. Brain Sciences, 7(6). 10.3390/brainsci7060058

Pylkkänen, L., Llinás, R., & Murphy, G. L. (2006). The representation of polysemy: MEG evidence. Journal of Cognitive Neuroscience, 18(1), 97–109. 10.1162/089892906775250003

Pylkkänen, L., Stringfellow, A., & Marantz, A. (2002). Neuromagnetic evidence for the timing of lexical activation: An MEG component sensitive to phonotactic probability but not to neighborhood density. Brain and Language, 81(1–3), 666–678. 10.1006/brln.2001.2555

Qin, L., & Gao, J. H. (2021). New avenues for functional neuroimaging: ultra-high field MRI and OPM-MEG. Psychoradiology, 1(4), 165–171. 10.1093/psyrad/kkab014

Salmelin, R. (2007). Clinical neurophysiology of language: The MEG approach. Clinical Neurophysiology, 118(2), 237–254. 10.1016/j.clinph.2006.07.316

Sander, T., Jodko-Władzińska, A., Hartwig, S., Brühl, R., & Middelmann, T. (2020). Optically pumped magnetometers enable a new level of biomagnetic measurements. Advanced Optical Technologies, 9(5), 247–251. 10.1515/aot-2020-0027

Sander, T., Preusser, J., Mhaskar, R., Kitching, J., Trahms, L., & Knappe, S. (2012). Magnetoencephalography with a chip-scale atomic magnetometer. 3(5), 27167–27172.

Seymour, R. A., Alexander, N., Mellor, S., Neill, G. C. O., Tierney, T. M., Barnes, G. R., & Maguire, E. A. (2021). NeuroImage Using OPMs to measure neural activity in standing, mobile participants. NeuroImage, 244(August), 118604. 10.1016/j.neuroimage.2021.118604

Seymour, R. A., Alexander, N., Mellor, S., O’Neill, G. C., Tierney, T. M., Barnes, G. R., & Maguire, E. A. (2022). Interference suppression techniques for OPM-based MEG: Opportunities and challenges. NeuroImage, 247(December 2021), 118834. 10.1016/j.neuroimage.2021.118834

Stockall, L., Stringfellow, A., & Marantz, A. (2004). The precise time course of lexical activation: MEG measurements of the effects of frequency, probability, and density in lexical decision. Brain and Language, 90(1–3), 88–94. 10.1016/S0093-934X(03)00422-X

Tan, G., Gai, J., Li, F., Huang, Q., Yu, W., Huang, Y., Zhang, G. Y., Zhao, X., Lin, Q., & Hu, Z. (2025). A Brain-Computer Interface System based on Magnetoencephalography with Optically Pumped Magnetometers. IEEE Transactions on Instrumentation and Measurement, PP, 1. 10.1109/TIM.2024.3522701

Tanner, D., Morgan-short, K., & Luck, S. J. (2015). How inappropriate high-pass filters can produce artifactual effects and incorrect conclusions in ERP studies of language and cognition. 52, 997–1009. 10.1111/psyp.12437

Tarkiainen, A., Helenius, P., & Salmelin, R. (2003). Category-specific occipitotemporal activation during face perception in dyslexic individuals: An MEG study. NeuroImage, 19(3), 1194–1204. 10.1016/S1053-8119(03)00161-7

Tierney, T. M., Alexander, N., Mellor, S., Holmes, N., Seymour, R., O’Neill, G. C., Maguire, E. A., & Barnes, G. R. (2021). Modelling optically pumped magnetometer interference in MEG as a spatially homogeneous magnetic field. NeuroImage, 244(July), 118484. 10.1016/j.neuroimage.2021.118484

Tsigka, S., Papadelis, C., Braun, C., & Miceli, G. (2014). Distinguishable neural correlates of verbs and nouns: A MEG study on homonyms. Neuropsychologia, 54(1), 87–97. 10.1016/j.neuropsychologia.2013.12.018

van Driel, J., Olivers, C. N. L., & Fahrenfort, J. J. (2021). High-pass filtering artifacts in multivariate classification of neural time series data. Journal of Neuroscience Methods, 352(November 2019), 109080. 10.1016/j.jneumeth.2021.109080

Wu, H., Liang, X., Wang, R., Ma, Y., Gao, Y., & Ning, X. (2024). A Multivariate analysis on evoked components of Chinese semantic congruity: an OP-MEG study with EEG. Cerebral Cortex, 34(4). 10.1093/cercor/bhae108

Xia, Q., & Peng, G. (2022). The roles of object and action, and concreteness and imageability, in the distinction between nouns and verbs: An ERP study on monosyllabic words in Chinese. Journal of Neurolinguistics, 61(August 2021), 101026. 10.1016/j.jneuroling.2021.101026

Xia, Q., Wang, L., & Peng, G. (2016). Nouns and verbs in Chinese are processed differently: Evidence from an ERP study on monosyllabic and disyllabic word processing. Journal of Neurolinguistics, 40, 66–78. 10.1016/j.jneuroling.2016.06.002

Xun, E., Rao, G., Xiao, X., & Zang, J. (2016). Dashuju beijing xia BCC yuliaoku de yanzhi [The construction of the BCC corpus in the age of Big Data]. Yuliaoku Yuyanxue [Corpora Linguistics], 3(1), 93–109.

Yu, X., Mancha, S., Tian, X., & Lau, E. (2024). Shared neural computations for syntactic and morphological structures: evidence from Mandarin Chinese. BioRxiv. 10.1101/2024.01.31.578104

Zhan, W., Guo, R., Chang, B., Chen, Y., & Chen, L. (2019). Beijing Daxue CCL Yuliaoku de Yanzhi [The building of the CCL corpus: its design and implementation]. Yuliaoku Yuyanxue [Corpus Linguistics], 6(1), 71–86.

Zhang, G., Garrett, D. R., & Luck, S. J. (2024a). Optimal filters for ERP research I : A general approach for selecting filter settings. November 2023. 10.1111/psyp.14531

Zhang, G., Garrett, D. R., & Luck, S. J. (2024b). Optimal filters for ERP research II : Recommended settings for seven common ERP components. January, 1–24. 10.1111/psyp.14530

Zhang, X., Chen, C., Zhang, M., Ma, C., Zhang, Y., Wang, H., Guo, Q., Hu, T., Liu, Z., Chang, Y., Hu, K., & Yang, X. (2020). Detection and analysis of MEG signals in occipital region with double-channel OPM sensors. Journal of Neuroscience Methods, 346(April), 108948. 10.1016/j.jneumeth.2020.108948

