## Supplementary Materials for "Unleashing the potential of OPM-MEG to study event-related fields against low-frequency artifacts: the case of sentence processing"

**N400 effects across the four conditions**. We performed a 2 × 2 (word type: nouns vs verbs; morpho-syntactic complexity: simpler vs. more complex) repeated-measures ANOVA on ERF for F7 across the 300-400 ms time window. When only employing a 0.1 Hz HPF, the main effect of word type was not significant, $F(1,10) = 0.05$, $p = 0.83$, partial $\eta^2 = 0.01$ (the main effect of morpho-syntactic complexity and the interaction effect *p's* > 0.76). When employing a 0.1 Hz HPF and DSS, the main effect of word type was also not significant, $F(1,10) = 1.94$, $p = 0.19$, partial $\eta^2 = 0.16$ (other effects *p's* > 0.63). When only employing a 1 Hz HPF, the main effect of word type was not significant as well, $F(1,10) = 2.77$, $p = 0.13$, partial $\eta^2 = 0.22$ (other effects *p's* > 0.54). When employing a 1 Hz HPF and DSS, the main effect of word type was statistically significant, $F(1,10) = 7.95$, $p = 0.018$, partial $\eta^2 = 0.44$ (other effects *p's* > 0.20). When only employing a 4 Hz HPF, the main effect of word type was not significant, $F(1,10) = 1.22$, $p = 0.30$, partial $\eta^2 = 0.11$ (other effects *p's* > 0.29). When employing a 4 Hz HPF and DSS, the main effect of word type was not significant as well, $F(1,10) = 0.79$, $p = 0.40$, partial $\eta^2 = 0.07$ (other effects *p's* > 0.30). See Figure S1.

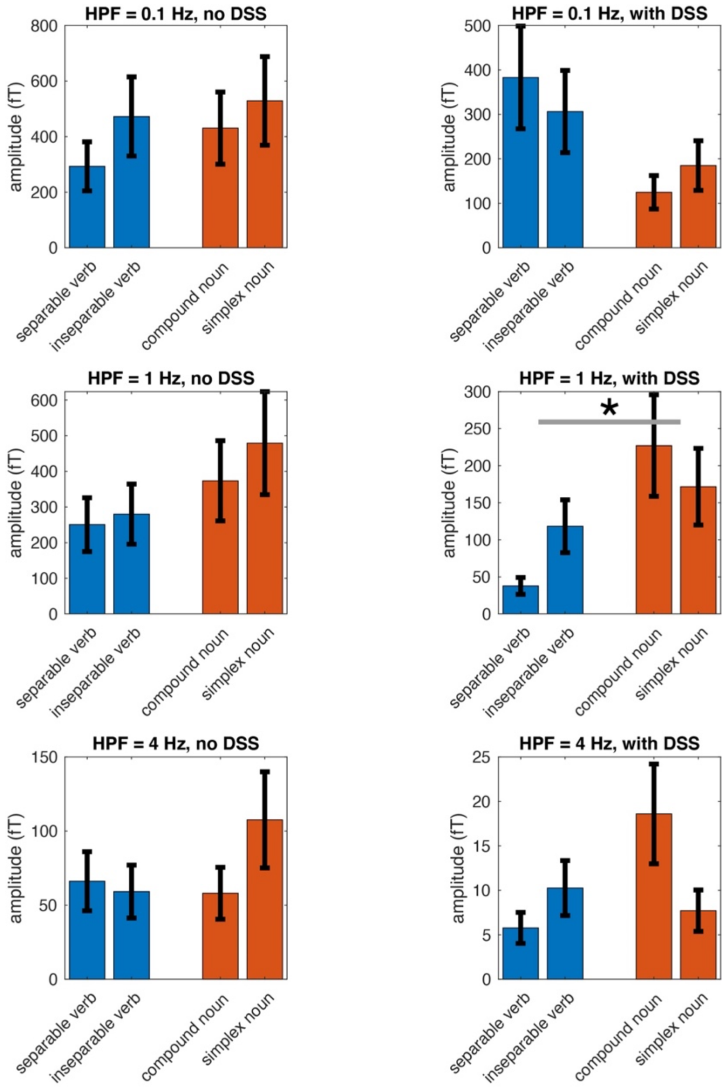


**Figure S1.** Comparison of the response amplitude at sensor F7 in the N400 time window (300-400 ms) across the four conditions. Only when an HPF (high-pass filter) of 1 Hz was combined with DSS, were we able to observe a significant main effect of word type (verb vs. noun). Error bars stand for standard error. *: $p < 0.05$. HPF: high-pass filter; DSS: denoising source separation.
